# Opposing effects of impulsivity and mindset on sources of science selfefficacy and STEM interest in adolescents

**DOI:** 10.1101/377994

**Authors:** L.K. Marriott, L. Coppola, Mitchell S.H., J. Bouwma-Gearhart, Z. Chen, D. Shifrer, J.S. Shannon

## Abstract

Impulsivity has been linked to academic performance in the context of Attention Deficit Hyperactivity Disorder, though its influence on a wider spectrum of students remains largely unexplored, particularly in the context of STEM learning (i.e. science, technology, engineering, and math). STEM learning was hypothesized to be more challenging for impulsive students, since it requires the practice and repetition of tasks as well as concerted attention to task performance. Impulsivity was assessed in a cross-sectional sample of 2,476 students in grades 6-12. Results show impulsivity affects a larger population of students, not limited to students with learning disabilities. Impulsivity was associated with lower sources of science self-efficacy (SSSE) scores, interest in all STEM domains (particularly math), and self-reported STEM skills. The large negative effect observed for impulsivity was opposed by “growth” mindset, which describes a student’s belief in the importance of effort when learning is difficult. Mindset had a large positive effect, which was associated with greater SSSE, STEM interest, and STEM skills. When modeled together, results suggest that mindset interventions may benefit impulsive students who struggle with STEM. Together, these data suggest important interconnected roles for impulsivity and mindset that can influence secondary students’ STEM trajectories.

## Introduction

Students’ self-beliefs about their abilities in STEM (i.e. science, technology, engineering, and math) directly correlate with persistence in STEM fields (1, 2), even independent of parents’ education or family income (3). The secondary school period is an important time for shaping students’ self-beliefs in STEM (3, 4) as well as for building STEM interest. While early interest in science is an important predictive factor for students later choosing a STEM-related career (5, 6), it can be over-shadowed by poor academic performance in math and science courses, thereby altering a student’s self-belief in their ability to succeed in science (3). These self-beliefs are thought to contribute to student attrition from STEM fields (5, 7).

Spinella (8) previously reported impulsivity to be negatively associated with academic grades in college-aged students. Impulsivity describes “a predisposition toward rapid, unplanned reactions to internal or external stimuli without regard to the negative consequences of these reactions to the impulsive individuals or to others” (9). More operationally, impulsivity describes two different behavioral characteristics: 1) an impairment of behavioral inhibition; and 2) a pronounced de-valuation of delayed outcomes (10, 11). Higher levels of impulsivity are associated with various psychopathologies including certain ADHD subtypes, substance use disorder, conduct disorder, and delinquency (12-16). In contrast, low impulsivity levels have been associated with compulsivity, obsessive compulsive disorder, and some eating disorders (17, 18). Thus, all individuals would be expected to fall along a continuous scale of impulsivity.

Most impulsivity research investigating academic performance has focused on the contexts of attention-deficit/hyperactivity disorder (ADHD) (19, 20), risky behaviors (21, 22), and early childhood self-control/regulation (23, 24) leaving the role of impulsivity as an underlying behavioral trait that may shape students’ academic performance largely unexplored (8, 25), particularly in the context of STEM learning. Impulsive students can have trouble staying on task and may be expected to find STEM learning more challenging, as academic effort in STEM involves practice and repetition of tasks as well as concerted attention to task performance. This may be especially true for mathematics, where content builds on prior knowledge and considerable repetitive practice is needed for mastery. For students, impulsivity may manifest as postponing homework or studying, which can contribute to poor academic performance. As students’ self-beliefs in STEM formed during secondary school can be negatively influenced by poor academic performance (3), it is possible that impulsivity may influence these relationships. For example, children diagnosed with ADHD can have trouble in school with sustained attention, hyperactivity, and impulsivity, which can negatively affect learning outcomes (26). Students with ADHD attain lower academic levels than their peers (27), an effect also found for children who are severely inattentive, hyperactive, and impulsive, but lack a formal diagnosis of the disorder (20, 28, 29). In the United States, the prevalence of these disorders among children and adolescents range from 5.9%-7.1% for ADHD (30), 5-6% for learning disabilities (31), and 0.6-2.2% for autism spectrum disorder (32). However, sub-clinical levels of impulsivity may also affect students with or without learning disability classifications.

This study explored the prevalence of impulsivity in a large cross-sectional sample of secondary students, when interest in science is being shaped (5, 6), to understand whether sub-clinical levels of impulsivity may affect a larger spectrum of students than previously considered. This study was not designed to be causal nor to identify learning disabilities among students, but rather to explore whether students’ impulsivity levels were associated with early measures of STEM persistence, such as STEM interest, science self-efficacy, and self-beliefs toward science and learning.

## Materials and Methods

### Participants and Settings

A total of six schools were recruited to participate in the current study with procedures overseen by OHSU Institutional Review Board (IRB, (protocol #3694). Schools had a prior academic relationship with the investigator (L.K.M.) and were recruited based on school sociodemographics. Schools were offered $500 USD for administering two anonymous surveys to their students during the 2014-2015 school year, with all sites accepting. Sites were distributed across three states (Oregon 1=two rural schools, 6-8th grades; Washington 2=one suburban school, 6^th^-8^th^ grades; California 3=three urban schools, one 7^th^-8^th^, one 9-12^th^, and one 7^th^-12th grades) (35). All sites permitted use of their facilities, managed interaction with students, and oversaw parental opt-out forms that maintained student anonymity to study staff.

### Assessment Procedures

Two paper-based surveys, approximately 30 minutes in length, were administered to students and separated by one month to lessen survey fatigue for students and class interruption time. To maintain anonymity while permitting the linkage of the two surveys, students were asked a series of questions on each survey to generate a unique identification number including a) first two letters of mother’s first name (ID_a), b) day of birth (ID_b), c) last two digits of phone number (ID_c), and d) birth order (ID_d). These responses, along with grade, gender, age, and teacher administering the survey, comprised the students’ “unique ID” and was used to match the two surveys using a deterministic matching procedure (described below).

### Instruments

Instruments included in Survey 1 included impulsivity, mindset, science self-efficacy, and STEM skills. Instruments included in Survey 2 assessed STEM domain interest, interest in a STEM career, and questions about learning behaviors.

- <u>Barratt Impulsiveness Scale</u> – short form (BIS-15) – The BIS-15 (36) comprised 15 items measured on a 4-point Likert scale (1-4, with six items reverse scored as previously reported (36, 37). Subscales (Attentional [A], Motor [M], and Non-Planning [NP]) previously produced Cronsbach’s alpha coefficients (α) between α=.60-.78 in university students. In the current study, a total of 2080 students completed all 15 items (α=.75), calculated from its three subscales, A (α=.74, n=2289), M (α=.61, n=2282), and NP (α=.68, n=2273).
- <u>Sources of Science Self-Efficacy</u> –SSSE applied Usher and Parajes’ validated mathematics scale (38) reworded for science (39). The instrument comprised 24 items that addressed four constructs: mastery experiences (ME), vicarious experiences (VE), social persuasion (P), and psychological and affective state (PH). Items were scored based on a 6-point Likert scale (0-5, scores from 0-120). Previous test reliability among 1225 middle and high school students produced α=.87, .71, 85, and .86 for the four constructs, respectively. In the current study, a total of 1899 students completed all 24 items (α=.86), representing a composite measure of SSSE calculated from ME (α=.88, n=2210), VE (α=.89, n=2145), P (α=.91, n=2086), and PH (α=.92, n=2088).
- <u>Mindset</u> – Mindset describes the continuum of a student’s felt beliefs of being able to increase personal intelligence through effort (termed “growth mindset”) versus it being a static trait conferred at birth (“fixed mindset”, 33, 34). A 20 item instrument designed by Dweck (33, 34) was scored on a 4-point Likert scale (1-4, with 10 items reverse-scored). Items stem from the Theory of Intelligence scale (33), Effort Belief Scale (40), and Patterns of Adaptive Learning Survey (41). Current analyses of 1759 students completing all 20 items produced α=.75.
- <u>STEM Skills</u> – Four questions assessed self-reported skills related to using and interpreting data. Each question offered the stem “I am good at projects involving...” with responses of 1) “using a website”; 2) “using data”; 3) “creating graphs”; and 4) “interpreting graphs”. Responses were scored on 5-point Likert scale ranging from Strongly Disagree to Strongly Agree. The current analyses of 2405 students completing the 4 items produced α=.76.
- <u>STEM Interest</u> – A 25-item STEM Semantics survey assessed student perceptions and interest across five STEM domains: 1) science, 2) math, 3) engineering, 4) technology, and 5) a STEM career (42, 43). Each domain included five questions that used adjective pairs to bookend a 7-point Likert scale, with a subset of items reverse scored. Domain scores were summed for each five question set. A composite STEM interest score was summed from all five subscales. Previous reliability among 174 students ranged from α=.84-.93, with 1575 students completing all 25 items in the current study (α=.93). The five subscales included science interest (α=.89, n=1807), math interest (α=.90, n=1812), engineering interest (α=.90, n=1755), technology interest (α=.90, n=1784), and interest in a STEM career (α=.92, n=1785).
- <u>STEM Learning</u> – Four questions from the Index of Learning Styles (44) were used to triangulate findings, as they dichotomize students’ processes for solving mathematics problems and overall learning pace in the context of impulsivity. Selected questions included: 1) “When I am doing long calculations: a) I tend to repeat all my steps and check my work carefully, or b) I find checking my work tiresome and I have to force myself to do it”; 2) “When I solve math problems: a) I usually work my way to the solutions one step at a time, or b) I often just see the solutions but then have to struggle to figure out the steps to get to them”; 3) “I learn: a) at a fairly regular pace. If I study hard, I’ll “get it”, or b) in fits and starts. I’ll be totally confused and then suddenly it all “clicks”“; and 4) “In a study group working on difficult material, I am more likely to a) jump in and contribute ideas, or b) sit back and listen”.

### Survey Processing and Statistical Analyses

Paper surveys were scanned using Remark software that populated survey data into Excel for statistical analyses by SAS and SPSS. Survey data were first matched in SAS 9.4 with subsequent analyses conducted with IBM SPSS Statistics, version 22. Geographical location and school demographics were obtained from 2013-2014 NCES data (35).

<u>Survey Linking Procedure.</u> A deterministic matching procedure was used to first match all nine variables (school, gender, grade, ID_a, ID_b, ID_c, age, teacher, and ID_d) with matched records moved to a new dataset. The procedure was repeated down to five variables, with handwriting samples confirming matching at each level (n=31 total; 100% agreement). This procedure was used to link Survey 1 (n=2476) with Survey 2 (n=2115), representing a conservative match rate of 41.4% (n=875) of anonymous students. Analyses were conducted on all completed items; therefore, comparisons within a survey had larger sample sizes than between surveys.

<u>Statistical Analyses.</u> Likert scale responses were converted numerically and summed for each subscale and composite score. Blank entries were not included in calculations. Non-parametric tests were first run on all comparisons (Mann Whitney U, Kruskal-Wallis H) due to controversy with Likert scale data (45, 46); however, no outcome differences were observed between any metrics using non-parametric versus parametric analyses, therefore, parametric tests were used for reporting in the current study. Results apply independent sample t tests and ANOVA using mean and standard deviation (SD). Bonferroni post-hoc tests were used to determine differences between groups through multiple comparisons. Scales were first analyzed as continuous variables before being binned into quartiles to support data visualization (e.g., impulsivity, mindset, SSSE, and math interest). Scale data were binned by quartile using SPSS and analyzed by general linear modeling to determine interactions between groups. Effect sizes are described using partial eta squared to account for comparisons across groups (47), with established benchmarks defining small (partial η^2^= 0.01), medium (partial η^2^ = 0.06), and large (partial η^2^= 0.14) effects (47, 48). Confidence intervals were calculated using SPSS syntax developed by Karl Wuensch (49). Finally, hierarchical linear modeling was used to account for the fact that multiple students from the same school may be more similar in responses than students at other schools. Specifically, the linear mixed model function in SPSS was used to account for parameter estimates of impulsivity, mindset, grade, gender and school on SSSE as fixed factors. Chi square tests were used to determine dichotomous differences in STEM learning across student quartiles. Graphs reflect mean+SEM, or percentages for chi square results.

<u>Missing Data Procedures.</u> To control for missing data, since impulsive students may be more likely to skip questions or scales, survey responses were analyzed by student demographics (gender and grade) within and between survey time points. Instrument scores were compared by completion status and demographics to understand if scores differed for students who completed all scales versus a subset of scales.

## Results

### Participants

A total of 3234 students were enrolled across the six sites (NCES 2015) and had the opportunity to complete survey measures, with 2476 completing Survey 1 and 2115 completing Survey 2. Fig 1 describes inclusion criteria and instrument sample sizes for analyses across the two survey time points. Of the 2476 students in grades 6-12 completed Survey 1, 85.8% were middle school students in U.S. grades 6-8 (Table 1). Participants were 47% female, consistent with NCES data for these participating schools (47.7% female; 58.7% qualify for free or reduced lunch). Racial/ethnic demographics of students were not collected in this study, though NCES data describe that 33.4% qualified as underrepresented minorities (URM) in STEM (50), denoting students who identified as African American (9.5%), Hispanic or Latino (22.6%), or Native American/Alaskan Native (1.3%). Students identifying as “Two or More Races” represent an additional 6.8% of the student population. Survey 2 was completed by a similar number of students, with chi square showing similar distributions in gender (p=.66) but not grade (p<0.001), as less 7^th^ grade students and more high school students participated in Survey 2 (Table 1).

**Fig 1.**
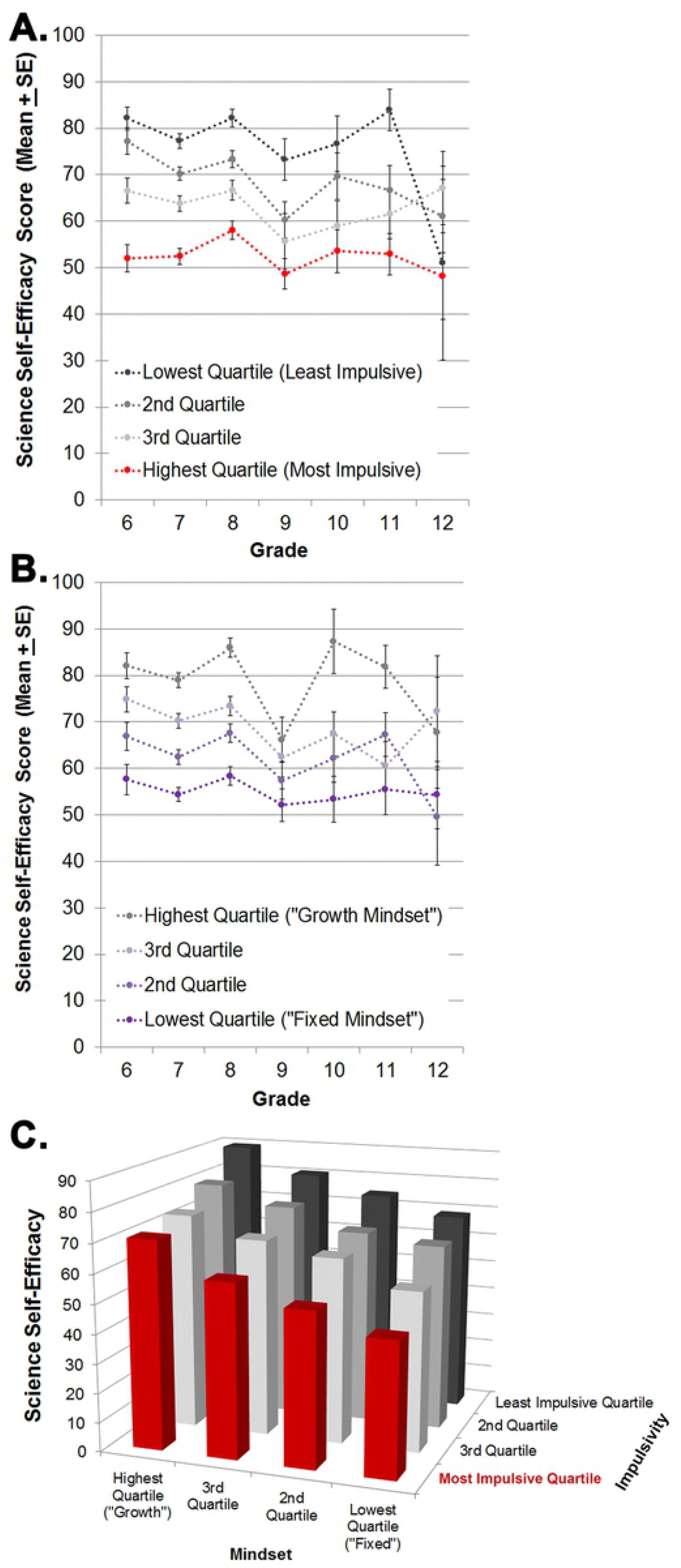
Inclusion criteria and sample sizes for analyses.

**Table 1.** Participant Demographics

|  | **Overall School Demographics**<br>NCES Data<br>(n=3234) | **Survey 1**<br>(n=2476)<br>*Mindset, Impulsivity,*<br>*Science Self-Efficacy,*<br>*STEM Skills* | **Survey 2**<br>(n=2115)<br>*STEM Interest, Learning*<br>*Behaviors* |
| --- | --- | --- | --- |
| **Gender** |  |  |  |
| Female | 1543 (47.7%) | 1132 (46.9%) | 972 (46.8%) |
| Male | 1691 (52.3%) | 1281 (53.1%) | 1103 (53.2%) |
|  |  | n=2413 | N=2075 |
| **Grade** |  |  |  |
| 6 | 493 (15.2%) | 449 (18.6%) | 422 (20.4%) |
| 7 | 1089 (33.7%) | 1011 (42.0%) | 658 (31.9%) |
| 8 | 1109 (34.3%) | 607 (25.2%) | 597 (28.9%) |
| 9 | 157 (4.9%) | 136 (5.6%) | 132 (6.4%) |
| 10 | 153 (4.7%) | 87 (3.6%) | 116 (5.6%) |
| 11 | 106 (3.3%) | 92 (3.8%) | 83 (4.0%) |
| 12 | 127 (3.9%) | 26 (1.1%) | 56 (2.7%) |
|  |  | n=2408 | n=2064 |

## Behavioral Measures

### Impulsivity

A total of 2080 students completed the impulsivity scale (mean=33.2, SD=6.7; Table 2). Quartiles denote scores of <= 28 (least impulsive); 29–33, 34–37, and 38+ (most impulsive), which were used to investigate relationships between mindset and STEM metrics (STEM interest and science self-efficacy). No differences in impulsivity subscales were observed for gender (subscale data not shown). Grade had a small effect on impulsivity (p<0.005; partial η^2^= 0.01), with similar effects observed for both M (p<0.001, partial η^2^= 0.013) and A (p<0.005; partial η^2^= 0.016) subscales. Specifically, 9^th^ graders had highest impulsivity as well as motor and attentional subscale scores, though differences were only significant when compared to 6^th^ grade students (p<0.05).

**Table 2.** Means and effect sizes of impulsivity, mindset, SSSE, and STEM domain interest across gender and grade.

|  | Impulsivity | Mindset | SSSE | Science Interest | Math Interest | Engineering Interest | Technology Interest | Interest in a STEM Career | Interest in STEM Domains (Cumulative Score) |
| --- | --- | --- | --- | --- | --- | --- | --- | --- | --- |
| Overall Mean + SD (n) | 33.2, 6.7, n=2080 | 60.0, 7.3, n=1759 | 67.4, 22.6, n=1899 | 24.2, 7.9 (n=1807) | 21.8, 8.8 (n=1812) | 22.6, 8.8 (n=1755) | 25.3, 8.2 (n=1784) | 24.0, 8.5 (n=1785) | 94.1, 24.2 (n=1575) |
| Gender | $t(2028)=1.20$, $p=0.23$<br>partial $\eta^2=0.001$ | $t(1723)=-1.99$, $p<0.05$<br>partial $\eta^2=0.002$ | $t(1856)=2.69$, $p=0.007$<br>partial $\eta^2=0.004$ | $t(1773)=2.76$, $p<0.007$<br>partial $\eta^2=0.004$ | $t(1781)=2.37$, $p<0.02$<br>partial $\eta^2=0.003$ | $t(1727)=12.05$, $p<0.001$<br>partial $\eta^2=0.078$ | $t(1754)=8.88$, $p<0.001$<br>partial $\eta^2=0.043$ | $t(1756)=5.8$, $p<0.001$<br>partial $\eta^2=0.019$ | $t(1773)=2.76$, $p<0.007$<br>partial $\eta^2=0.049$ |
| Male | 33.3, 6.4, n=1090 | 59.7, 7.5, n=896 | 68.8, 21.9, n=967 | 24.7, 7.8, n=939 | 22.3, 8.6, n=945 | 24.9, 8.4, n=918 | 26.9, 8.0, n=928 | 25.2, 8.4, n=926 | 99.1, 23.7, n=825 |
| Female | 32.9, 6.8, n=940 | 60.4, 7.0, n=829 | 66.6, 23.3, n=891 | 23.7, 8.0, n=836 | 21.26, 9.0, n=838 | 20.0, 8.6, n=811 | 23.5, 8.0, n=828 | 22.8, 8.5, n=832 | 88.3, 23.6, n=726 |
| Grade | $F(6, 2022)=3.32$, $p<0.004$<br>partial $\eta^2=0.01$ | $F(6, 1719)=2.47$, $p<0.03$<br>partial $\eta^2=0.009$ | $F(6, 1850)=7.31$, $p<0.001$<br>partial $\eta^2=0.023$ | $F(6, 2148)=11.56$, $p<0.001$<br>partial $\eta^2=0.022$ | $F(6, 2086)=10.36$, $p<0.001$<br>partial $\eta^2=0.029$ | $F(6, 2040)=3.76$, $p<0.001$<br>partial $\eta^2=0.018$ | $F(6, 2034)=4.24$, $p<0.001$<br>partial $\eta^2=0.019$ | $F(6, 1863)=7.93$, $p<0.001$<br>partial $\eta^2=0.038$ | $F(6, 2148)=11.56$, $p<0.001$<br>partial $\eta^2=0.031$ |
| 6 | 32.0, 6.6, n=371<sup>b</sup> | 61.3, 6.8, n=218<sup>c</sup> | 71.3, 23.1, n=268<sup>a</sup> | 25.5, 7.5, n=352<sup>a</sup> | 24.1, 8.8, n=355<sup>a</sup> | 23.4, 8.3, n=329<sup>a</sup> | 26.5, 7.9, n=341<sup>a</sup> | 26.4, 7.8, n=347<sup>a</sup> | 98.4, 23.0, n=303<sup>a</sup> |
| 7 | 33.1, 6.3, n=847 | 59.9, 7.2, n=771 | 66.4, 21.7, n=797<sup>bnz</sup> | 24.1, 7.7, n=552<sup>a</sup> | 21.8, 8.5, n=556<sup>y</sup> | 23.2, 9.0, n=543<sup>a</sup> | 25.9, 8.2, n=542<sup>bz</sup> | 24.3, 8.5, n=547<sup>bz</sup> | 95.5, 23.2, n=479<sup>a</sup> |
| 8 | 33.3, 7.0, n=514 | 60, 7.4, n=464 | 70.4, 22.5, n=498<sup>a</sup> | 24.4, 8.1, n=516<sup>a</sup> | 21.8, 8.5, n=517<sup>y</sup> | 22.5, 8.8, n=505<sup>b</sup> | 24.7, 8.4, n=519 | 24.1, 8.2, n=513<sup>bx</sup> | 93.9, 24.5, n=448<sup>a</sup> |
| 9 | 34.6, 6.8, n=115<sup>y</sup> | 58.5, 7.1, n=111<sup>z</sup> | 58.3, 24.0, n=119<sup>mx</sup> | 20.4, 8.3, n=117 | 20.3, 8.9, n=118<sup>x</sup> | 19.2, 9.1, n=117 | 22.7, 8.0, n=116 | 21.0, 8.7, n=115<sup>x</sup> | 82.8, 25.0, n=106 |
| 10 | 34.3, 6.1, n=77 | 59, 7.2, n=68 | 64.0, 25.0, n=71 | 24.9, 7.3, n=101<sup>a</sup> | 20.2, 9.2, n=98<sup>x</sup> | 23.5, 8.6, n=100<sup>b</sup> | 26.7, 7.5, n=99<sup>b</sup> | 23.5, 9.5, n=102<sup>c</sup> | 95.9, 25.1, n=91<sup>b</sup> |
| 11 | 33.6, 7.2, n=83 | 60.5, 8, n=76 | 66.1, 23.9, n=81 | 23.5, 7.7, n=72 | 18.2, 8.8, n=72<sup>x</sup> | 21.1, 7.8, n=66 | 24.4, 7.3, n=73 | 20.8, 7.7, n=72<sup>x</sup> | 87.1, 23.0, n=60<sup>z</sup> |
| 12 | 34.1, 6.1, n=22 | 58.4, 7.3, n=18 | 57.9, 17.0, n=23 | 23.5, 8.8, n=51 | 18.4, 9.4, n=51<sup>x</sup> | 20.0, 8.7, n=52 | 22.9, 8.2, n=50 | 19.6, 9.2, n=51<sup>x</sup> | 84.5, 25.5, n=48<sup>y</sup> |

Results shown as Mean, SD, and sample size of analysis. Effect size benchmarks define small (partial $\eta^2=0.01$ ), medium (partial $\eta^2=0.06$ ), and large (partial $\eta^2=0.14$ ) effects. Bonferroni post-hoc tests were used to determine differences between groups through multiple comparisons. For grade, <sup>a</sup> denotes differences between 9<sup>th</sup> grade students at the $p<0.001$ , <sup>b</sup> $p<0.01$ , and <sup>c</sup> $p<0.05$ levels whereas <sup>x</sup> denotes differences between 6<sup>th</sup> grade students at the $p<0.001$ , <sup>y</sup> $p<0.01$ , and <sup>z</sup> $p<0.01$ levels.

### Mindset

A total of 1759 students completed the mindset instrument (mean=60.0, SD=7.3). Mindset quartiles reflect scores of <= 55 (lowest mindset, referred to in the literature as “fixed” mindset), 56 – 60, 61 – 65, to 66+ (highest mindset, “growth” mindset). Mindset scores were higher among females than males (p<0.05, Table 2), though the effect size was very small (partial η^2^= 0.002). A small but significant difference was observed across grade (p<0.03; partial η^2^= 0.009), relating to lower mindset scores among 9^th^ grade students compared to 6^th^ graders.

### Sources of Science Self-Efficacy (SSSE)

A total of 1912 students (mean=68.0, SD=22.6, Table 2) completed the SSSE scale with quartiles reflecting scores of <= 52, 53–67, 68–84, and 85+. SSSE scores were higher among males than females (p<0.001), though only the physiological state (PH) subscale differed between gender (p<0.001; partial η^2^= 0.013), with males having higher sub-scores than females (subscale data not shown). As PH items are reverse-scored, lower numbers denote a higher physiological response. Grade had a small effect on SSSE (p<0.001, partial η^2^= 0.023) with Bonferroni post-hoc tests showing lower SSSE and ME sub-scores among 9^th^ graders compared to students in 6-8^th^ grade (p<0.002).

### STEM Skills

A composite STEM skills score was calculated for 2405 students (mean=14.5, SD=3.1) from four questions (mean, SD, n) that asked about self-reported skills using a website (4.0, 0.9, n=2417), using data (3.7, 1.0, n=2411), creating graphs (3.5, 1.1, n=2412), and interpreting graphs (3.3, 1.1, n=2408). Females had significantly lower scores than males on all questions and the composite score (p<0.001; partial η^2^= 0.01) though no differences were found between grades (p=0.35).

### Interest in STEM Domains and Career Interest

Interest in all four STEM domains were quantified for 1575 students using Survey 2 responses (Table 2). STEM domain scores differed significantly between males and females (all p<0.02), with males having higher scores in each category. Effect sizes for gender ranged from small to medium and all STEM domains differed significantly by grade (Table 2, p<0.001), with small to small-medium effect sizes observed. Bonferroni tests showed 9^th^ graders had significantly lower interest across all domains.

### Impulsivity and Mindset have Opposing Effects on Sources of Science Self-Efficacy

Pearson product-moment correlations were first used to determine relationships between impulsivity, mindset, and sources of science self-efficacy among students in grades 6-12, with results consistent across all school sites and grades. Impulsivity was negatively associated with SSSE (r=-.43, n=1663, p<0.001), whereas mindset was positively associated (r=.40, n=1580, p<0.001). A series of two-way ANOVAs were conducted by general linear modeling (GLM) to examine the relationship between impulsivity and mindset quartiles on SSSE. Students in the least impulsive quartile (≤28) had highest mean SSSE scores (79.5±21.8), with SSSE scores declining significantly with each impulsivity quartile (Fig 2A; 71.0±20.6, 64.3±19.1, 53.7±21.8), resulting in a large effect size (*F*_(3, 1662)_ = 114.11, p<0.001, η^2^_*p*_ = 0.171, 90% CI [0.144, 0.196]). Post-hoc tests revealed SSSE scores differed between each impulsivity quartile. The significant effects of impulsivity on SSSE persisted when controlling for each M, A, and NP subscale scores (all p<0.001). Likewise, the effect was consistent across grade and no interaction was observed among 1673 students (p=0.85).

**Fig 2.**
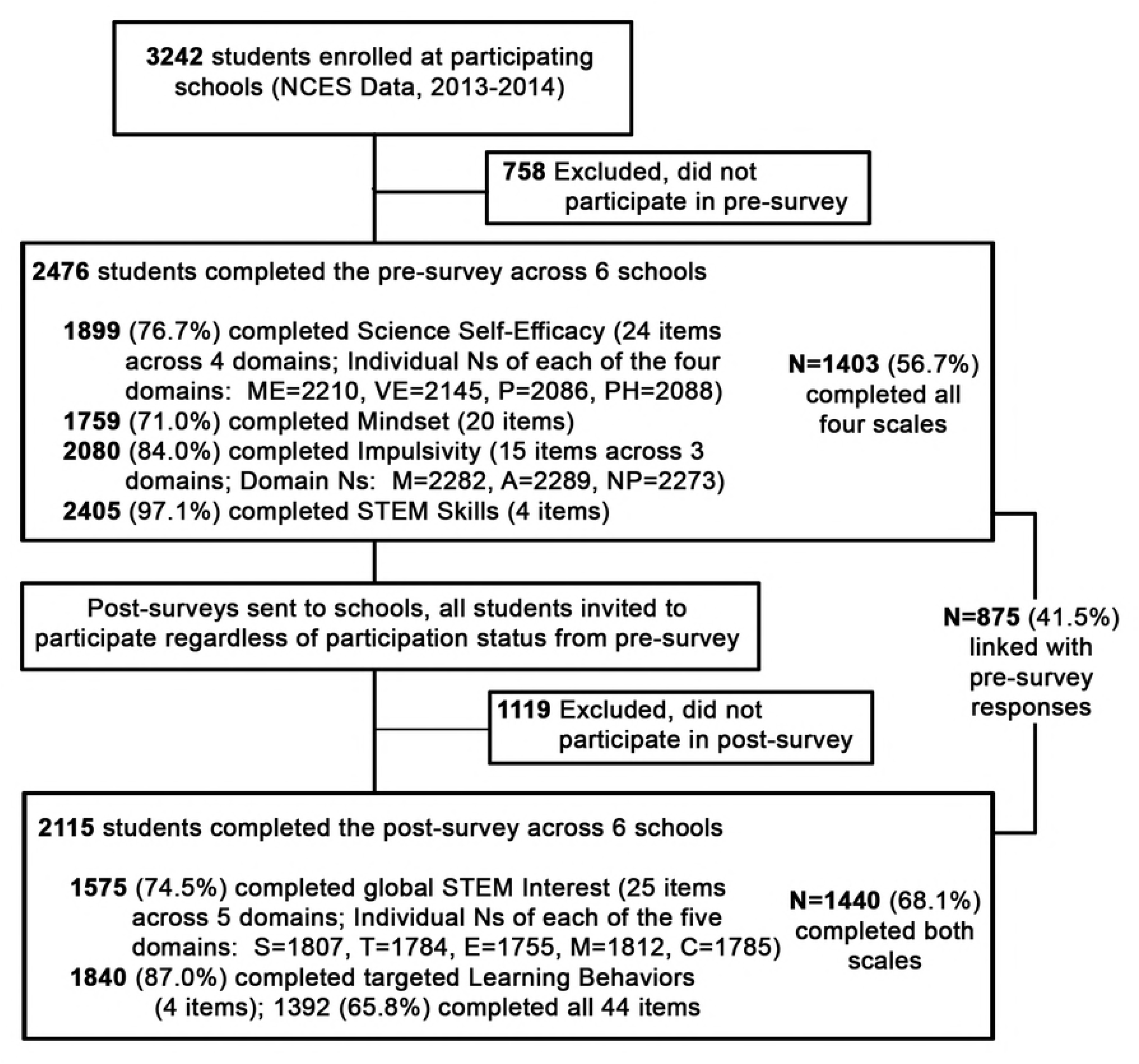
Sources of science self-efficacy scores were influenced by impulsivity. (A; large negative effect size; p<0.001; partial η^2^=0.171) and mindset (B; large positive effect size, p<0.001; partial η^2^=0.170). When modeled together (C), mindset opposed impulsivity’s negative stepwise effects on SSSE. Students with most impulsivity (red bars) yet highest mindset had equivalent science self-efficacy scores to students with least impulsivity yet lowest mindset.

Mindset positively associated with SSSE (n=1580, p<0.001), resulting in a large effect size (F_(3,1579)_ = 105.7, p<0.001, η^2^_*p*_ = 0.167, 90% CI [0.142, 0.196], Table 3). Mindset quartiles differed significantly, with students in the highest mindset quartile (66+) having higher mean SSSE scores (80.8+19.4) than students in lower mindset quartiles (Fig 2B; 70.8±22.0, 64.2±19.9, and 55.5±21.2). Grade affected mindset (p<0.001; partial η^2^= 0.009), where 9^th^ grade students had lower mindset scores than 6^th^, 7^th^, and 8^th^ grade students (all p<0.01), though no interaction was observed between mindset quartile and grade on SSSE (p=.79).

**Table 3.** Effect Sizes and correlation coefficients of impulsivity and mindset on STEM metrics,

| Metrics | Lowest Quartile<br>(Mean, SD, n) | Highest Quartile<br>(Mean, SD, n) | SS | Df, n | MS | F | Sig (p) | Effect Size<br>(Partial $\eta^2$) | 90% CI | r |
| --- | --- | --- | --- | --- | --- | --- | --- | --- | --- | --- |
| **Impulsivity on:** |  |  |  |  |  |  |  |  |  |  |
| Sources of Science Self-Efficacy | 79.5, 21.8, 432 | 53.7, 20.6, 406 | 149257.8 | 3, 1662 | 49752.6 | 114.11 | .000 | .171 | 0.144, 0.196 | -.43 |
| Composite STEM Domains Score | 101.9, 23.2, 149 | 81.9, 22.3, 131 | 28372.6 | 3, 566 | 9457.5 | 18.71 | .000 | .091 | 0.053, 0.126 | -.32 |
| Composite STEM Skills | 15.6, 2.9, 521 | 14.4, 3.1, 2064 | 1542.8 | 3, 2064 | 514.3 | 57.9 | .000 | .078 | 0.059, 0.096 | -.31 |
| Mathematics Interest | 25.0, 8.5, 164 | 18.9, 8.1, 149 | 3156.8 | 3, 642 | 1052.3 | 14.47 | .000 | .064 | 0.034, 0.093 | -.29 |
| Science Interest | 25.9, 7.6, 165 | 21.2, 7.6, 150 | 1919.0 | 3, 647 | 639.3 | 10.82 | .000 | .048 | 0.022, 0.074 | -.24 |
| Interest in a STEM Career | 25.7, 8.2, 164 | 21.6, 8.4, 144 | 1414.41 | 3, 630 | 471.5 | 6.68 | .000 | .031 | 0.01, 0.053 | -.18 |
| Engineering Interest | 24.2, 8.1, 161 | 19.9, 8.7, 145 | 1390.9 | 3, 624 | 463.6 | 6.42 | .000 | .030 | 0.009, 0.052 | -.19 |
| Technology Interest | 26.7, 7.5, 164 | 22.8, 8.7, 144 | 1273.0 | 3, 631 | 424.3 | 6.34 | .000 | .029 | 0.009, 0.051 | -.16 |
| **Mindset on:** |  |  |  |  |  |  |  |  |  |  |
| Sources of Science Self-Efficacy | 56.2, 21.4, 422 | 81.7, 19.3, 381 | 138994.5 | 3, 1588 | 46331.5 | 108.19 | .000 | .170 | 0.142, 0.196 | .41 |
| Composite STEM Domains Score | 85.8, 22.0, 117 | 103.2, 22.3, 118 | 18905.7 | 3, 472 | 6301.9 | 12.30 | .000 | .073 | 0.036, 0.108 | .25 |
| Science Interest | 21.6, 7.6, 139 | 26.8, 8.0, 134 | 1932.6 | 3, 530 | 644.2 | 10.71 | .000 | .058 | 0.026, 0.088 | .23 |
| Composite STEM Skills | 13.8, 3.6, 483 | 15.6, 2.9, 413 | 761.3 | 3, 1750 | 253.8 | 26.6 | .000 | .044 | 0.028, 0.059 | .22 |
| Interest in a STEM Career | 22.3, 8.1, 130 | 26.2, 8.4, 129 | 1478.5 | 3, 518 | 492.8 | 6.87 | .000 | .039 | 0.013, 0.065 | .17 |
| Technology Interest | 23.3, 8.8, 131 | 27.6, 6.8, 131 | 1299.8 | 3, 520 | 433.3 | 6.79 | .000 | .038 | 0.012, 0.064 | .18 |
| Mathematics Interest | 20.7, 8.8, 137 | 24.0, 9.0, 130 | 908.2 | 3, 528 | 302.7 | 3.97 | .008 | .022 | 0.003, 0.043 | .15 |
| Engineering Interest | 21.1, 8.9, 130 | 24.1, 8.4, 128 | 739.5 | 3, 513 | 246.5 | 3.19 | .024 | .018 | 0.001, 0.037 | .11<sup>a</sup> |

Items ranked by quartile effect size (partial $\eta^2$ ) for both impulsivity and mindset using established benchmarks to define small (partial $\eta^2$ = 0.01), medium (partial $\eta^2$ = 0.06), and large (partial $\eta^2$ = 0.14) effects (47, 48).

The combined effect of mindset and impulsivity on SSSE was examined by two-way ANOVA among 1405 students. Significant, stepwise effects in opposing directions were observed for both impulsivity and mindset quartiles on SSSE (all p<0.001; Fig 2C). Thus, mean SSSE scores for students in the most impulsive quartile/highest mindset quartile (70.8±2.9; 95% CI=65.2-76.5) were equivalent to students in the least impulsive/lowest mindset quartile (68.7±2.8 SE; 95% CI 63.3-74.1). These patterns were consistent within each middle school grade (6th-8th), which comprised >85% of the sample, and were reproducible for high school when collapsing grades 9-12, which comprised a smaller sample size. No interaction was observed between mindset and impulsivity on SSSE (p=0.71). Table 4 describes parameter estimates for SSSE using hierarchical linear modeling.

**Table 4.** Linear mixed model estimates for the fixed effects of impulsivity and mindset on sources of science self-efficacy.

| Parameters | Estimate | Standard Error | df | t | Significance (p) | 95% Confidence Interval |
| --- | --- | --- | --- | --- | --- | --- |
| Intercept | 287.225 | 76.381 | 1349 | 3.76 | .000 | 137.39, 437.06 |
| Grade | -25.839 | 9.658 | 1349 | -2.68 | .008 | -44.79, -6.89 |
| Impulsivity | -2.298 | 0.614 | 1349 | -3.74 | .000 | -3.50, -1.09 |
| Mindset | -2.401 | 1.183 | 1349 | -2.03 | .043 | -4.72, -0.08 |
| School | -0.784 | 0.324 | 1349 | -2.42 | .016 | -1.42, -0.15 |
| Gender | -3.409 | 5.438 | 1349 | -0.63 | .531 | -14.08, 7.26 |
| Gender * Impulsivity | -4.136 | 1.381 | 1349 | -2.99 | .003 | -6.85, -1.43 |
| Grade * Mindset | 0.374 | 0.151 | 1349 | 2.47 | .013 | 0.08, 0.67 |
| Grade * Impulsivity | 0.142 | 0.072 | 1349 | 1.98 | .047 | 0.00, 0.28 |
| Gender * Grade * Impulsivity | 0.452 | 0.175 | 1349 | 2.59 | .010 | 0.11, 0.80 |

Schwarz's Bayesian Criterion (BIC) score was 11936.8 and coefficients of variation were 33.8% (SSSE), 12.0% (Mindset), 20.0% (Impulsivity), 16.7% (grade), 33.8% (gender), and 51.8% (school).

### STEM Interest is positively associated with SSSE

Moderate, positive correlations were observed between science interest and SSSE (r=.48) and a large effect was observed (p<0.001, n=586, partial η^2^= 0.197). Interest in all STEM domains correlated with SSSE (all p<0.001), with strongest associations and largest effect sizes observed for composite STEM domain interest (r=.43, n=518, p<0.001, partial η^2^= 0.171) and STEM career interest (r=.32, n=583, p<0.001, partial η^2^= 0.103). Small-medium effect sizes were observed for all other STEM domain quartiles on SSSE (partial η^2^= 0.037-0.064). A series of two-way ANOVAs were conducted by hierarchical linear modeling to examine the relationship between mindset and impulsivity quartiles on each STEM interest domain (all p<0.001), with small to medium effect sizes observed for each measure (**Table 3**).

### SSSE is positively associated with students’ beliefs in their STEM skills

GLM revealed a large effect size of SSSE quartiles on STEM skills (*F*_(3, 1888)_ = 108.9, p<0.001, η^2^_*p*_ = 0.148, 90% CI [0.123, 0.171]), where students in the lowest SSSE quartile had significantly lower STEM skill scores (mean=12.9, SD=3.3, n=474) than students in the highest SSSE quartile (mean=16.2, SD=2.6, n=450). Both impulsivity and mindset influenced STEM skills (partial η^2^ =.078 and .044, respectively; Table 3), particularly graph interpretation, which had the lowest mean of all four questions.

### Effect of Gender on the Relationship between STEM metrics

Females had lower scores than males in composite STEM interest (*F*(6, 1516)=2.3, p<0.04, partial η^2^= 0.009) and math interest (*F*(6, 1745)=2.2, p<0.05; partial η^2^= 0.007), particularly in 9^th^ grade (p<0.05). Consistent with STEM Interest, females showed lower SSSE scores than males (p<0.001) and a significant stepwise relationship was observed between SSSE quartiles and composite STEM domain interest (p<0.001) that resulted in a large effect size (partial η^2^= 0.152). Specifically, females had lower composite STEM domain interest scores across all SSSE quartiles (partial η^2^= 0.005). Females also had lower SSSE scores than males across the lowest three mindset quartiles, though equivalent scores were observed between genders in the highest mindset quartile (*F*(3,1548)=3.1, p<0.05; partial η^2^= 0.004). No interactions were observed for science (p=.24), engineering (p=.17), technology (p=.51), or STEM careers (p=.09).

### Conserved Relationship between Impulsivity, Mindset, SSSE, and Math Interest

Math interest quartiles were calculated for students (lowest=≤16; 17-21; 22-29; 30+) to permit analyses of self-reported learning behaviors by chi square. Two questions asked students about their procedures when solving math problems, one asked about learning pace, and one asked about behaviors when working in a group setting. A striking pattern emerged across all four questions between high/low quartiles of students, where most impulsive students showed similar responses to students with least mindset, least SSSE, and least math interest (Fig 3).

**Figure 3:**
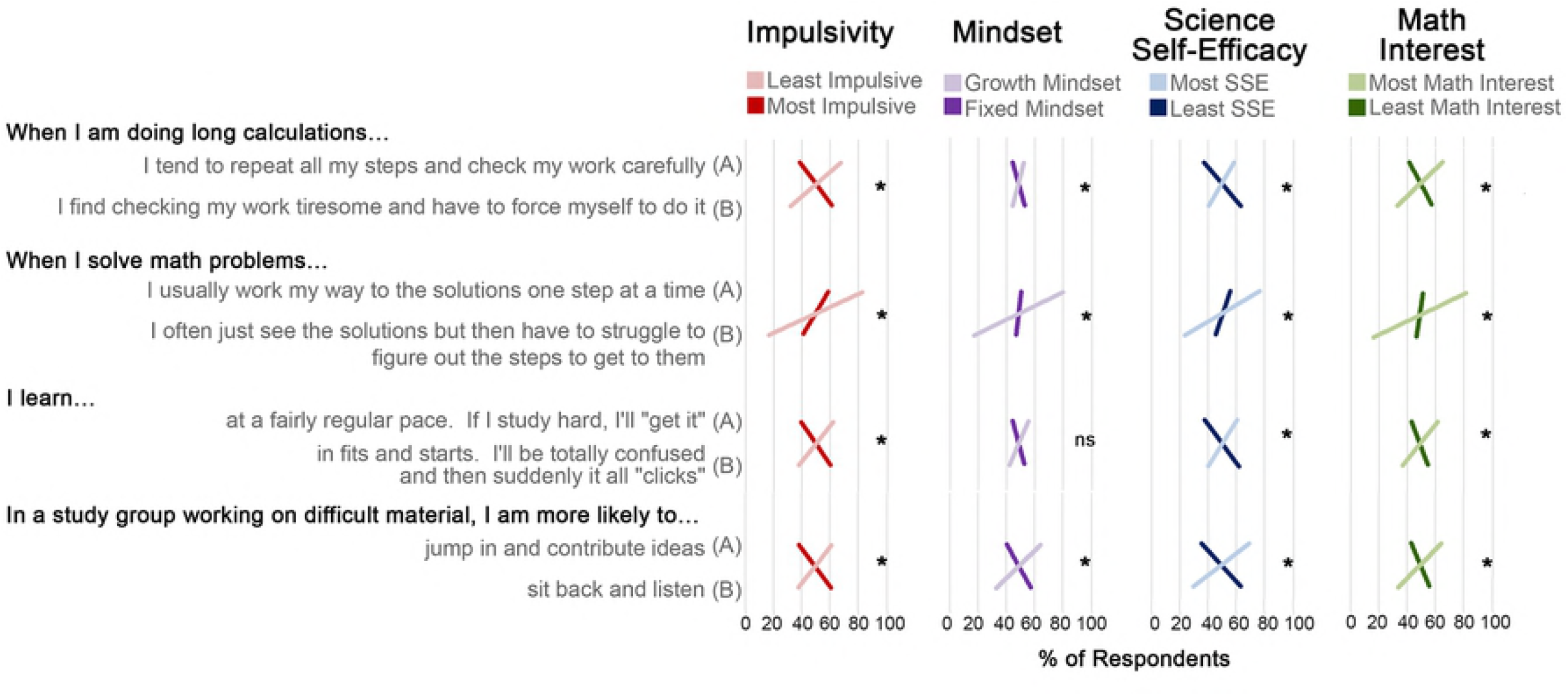
Learning behaviors are conserved when comparing highest quartiles of impulsivity with lowest quartiles of mindset, SSSE, and math interest. Most impulsive students (red lines) reported similar difficulties when solving math problems as students in lowest mindset, SSSE, and math interest quartiles (darker lines, *p<0.001).

### Missing Data Comparisons

Patterns of missing data were analyzed by instrument completion status, student demographics, and survey time points. A total of 1403 students (56.7%) completed all four Survey 1 scales (impulsivity, mindset, SSSE, and STEM skills), 734 (29.6%) partially completed scales (i.e., attempted all four scales, but did not fully complete at least one scale), 291 (11.8%) skipping at least one scale in entirety, and 48 students (1.9%) did not respond to any questions on all four scales. Completion status did not differ for students completing Survey 1 versus matched surveys (p=.90). Completion status was not affected by gender (p=.21), though grade had a significant effect (p<0.001) where 6^th^ grade students were less likely to complete all scales (37.4%) compared to other grades (59.9-69.6%). Rather than partial completions, 6^th^ grade students skipped entire scales (31.6%) compared to older students (2.2%-10.3%), possibly due to lack of time. No differences existed in instrument scores between Survey 1, Survey 2, or matched surveys for any of the instruments except SSSE, which was higher among students with matched surveys (M=68.9, SD=23.0, n=671) compared to Survey 1 alone (M=66.6, SD=22.4, n=1228; p<0.05). Students with partial or skipped instruments were grouped for analyses, though only mindset showed a significant difference based on completion status (p<0.01), with higher mindset scores among students completing all scales (60.2, SD 7.3, n=1403) than partial completions (M=59.0, SD=7.2, n=358). GLM was used to determine if differences existed in scores by completion status and demographics. When examined by completion status, only STEM career interest scores differed, where higher scores were observed among students with partial completions on other scales (p<0.05).

## Discussion

The research presented above confirms the positive association and large effect size between science self-efficacy and STEM domain interest demonstrated by others (3, 5, 6). It also confirms a positive association between “growth” mindset and self-beliefs towards STEM (51), which this study expands to include science self-efficacy (large effect size), interest in all STEM domains (small to moderate effect size), interest in a STEM career (small-moderate effect size), and self-beliefs in STEM skills, such as using data and interpreting graphs (moderate effect size) among students in grades 6-12. Consistent with previous findings showing impulsivity affecting academic performance in the context of ADHD and self-discipline (20, 24, 29), this manuscript reports a negative association of impulsivity on all measures of STEM studied, including sources of science self-efficacy (large effect size), interest in all STEM domains (small to moderate effect size), interest in a STEM career (small-moderate effect size), and STEM skills (moderate effect size). These findings suggest that impulsivity is likely influencing STEM learning outside the context of diagnosed and undiagnosed ADHD, which is estimated to have a prevalence within the U.S. school population of 5.9%-7.1% (30), though up to 11% per parent self-report (52). The data presented here offer that students fall along a continuum of impulsivity scores, with a negative stepwise effect observed for each impulsivity quartile on all STEM outcomes measured across a large, three state sample of adolescents in grades 6-12 (Table 3). Thus, while some students may have diagnosed or undiagnosed ADHD, these data support a larger reach of impulsivity that may negatively impact STEM persistence, possibly by influencing students’ self-beliefs in their STEM abilities.

These results are not designed to be causal, but rather offer preliminary support for the combined impact that the degree of impulsivity and growth mindset play as significant behavioral correlates of STEM interest and science self-efficacy (Fig 2C). For example, students in the most impulsive/highest mindset group had identical sources of science selfefficacy (SSSE) scores to students in the least impulsive/lowest mindset group. As impulsivity is thought to be a stable trait, whereas mindset can be grown, these findings suggest that mindset interventions may be beneficial for improving impulsive students’ self-efficacy for science. Growth mindset interventions, which emphasize recognition for effort rather than achievement, have been shown to improve learning and achievement (51, 53-55), particularly among groups underrepresented in STEM domains (40, 56-60). This may be particularly important, since currently, no classroom strategies have sufficient evidence for supporting learning gains among ADHD students, even following medication to alleviate symptoms (61, 62). This research suggests potential for mindset interventions, especially for students with highest impulsivity, and with respect to science and math, most notably.

These findings are supported by data describing similar patterns for how most impulsive students solve math problems and engage in learning (Fig 3), which mirror patterns observed for students with least mindset, least science self-efficacy, and least math interest. These cross-sectional findings offer that impulsive students may struggle more when solving math problems or learning difficult material, which may negatively influence self-beliefs in their abilities, consistent with previous reports (3). Impulsive students are not at an academic disadvantage, as their ability to perceive situations differently and learn at a different pace may be an asset in some situations, as early literature supports the notion that impulsivity can have functional or dysfunctional effects (63). For example, Tymms and Merrell (20) offer that blurting out answers may be an overt sign of cognitive engagement, where impulsivity may serve a positive function. Our data show that “when in a study group working on difficult material”, impulsive students were more likely to “sit back and listen” than “jump in and contribute ideas”. While seemingly counterintuitive, this finding may stem from impulsive students’ altered self-beliefs in their abilities when working on material that is challenging. For example, when restricting analyses to only the most impulsive quartile, students who “jump in and contribute ideas” had significantly higher sources of science self-efficacy scores (p<0.02), mastery experience sub-scores (p<0.01), science interest scores (p<0.05), and reported greater self-beliefs in their ability to interpret graphs (p<0.05) than equally impulsive students who reported to “sit back and listen”. No differences were observed for math interest (p=0.07) or mindset (p=.13) between these students. Thus, opportunities may exist for supporting impulsive students in STEM as they engage in difficult material or problem-based learning.

Consistent with prior studies documenting a gender gap in STEM (51, 64-66), this study observed females had lower sources of science self-efficacy, which confirm results from Britner and Parajes (67) using the same scale. This effect was not related to impulsivity, as no difference in impulsivity was observed between gender. However, mindset may play a role, as males had higher sources of science self-efficacy scores than females in the lowest three quartiles of mindset, despite equivalent scores in the highest mindset quartile (‘growth mindset’). Thus, targeting females for mindset interventions may be particularly successful if females’ self-beliefs toward their STEM abilities are low. Likewise, mindset interventions may also help students who express interest in STEM but lack the background content knowledge in a STEM domain, making the work more challenging, albeit surmountable. When not prepared for academic difficulties, students’ self-beliefs in their abilities may be challenged (56, 57) and reduce STEM interest and engagement (3). Finally, consistent with prior findings (68), 9^th^ grade students had lower sources of science self-efficacy, interest in STEM domains, interest in a STEM career, and mindset, as well as a slight but significant increase in impulsivity when compared to students in other grades. Given that 9^th^ grade is the time when students are told that their grades are first starting to ‘count’ towards college, students may feel greater stress to succeed academically and may decline STEM electives, particularly if grades are low and/or a student feels behind compared to peers.

Important limitations of this work relate to its lack of causal design as well as caution in interpretations for grades 10-12. While 12^th^ grade students also have low sources of science self-efficacy scores, the smaller sample size limits confidence in making interpretations related to effects of gender, mindset, or impulsivity. Instead, efforts focus primarily on middle school grades and have grouped high school grades 9-12 together prior to testing associations. In addition, the cross-sectional design separated surveys across two time points to ease survey fatigue, which resulted in a lower sample size when comparing relationships with STEM domain interest. While significant, greatest confidence can be attributed to relationships between impulsivity, mindset, and sources of science self-efficacy, as these measures were completed within the same survey and were highly reproducible in every school site studied. While a tendency for impulsive students to not complete a questionnaire was expected, this was not the case, as only mindset scores differed between students who completed all instruments versus students with partially completed or completely skipped instruments. Instead, 6^th^ grade students had the greatest amount of skipped instruments, rather than partial completions, likely due to survey length and limited time.

## Conclusion

This study offers that impulsivity may affect learning behaviors and self-beliefs regarding STEM across a wider spectrum of adolescents than previously considered. Based on the data, it is hypothesized that STEM persistence and attrition may be attributable to students’ underexplored behavioral characteristics (e.g., impulsivity and mindset) that reinforce or impede STEM learning, consistent with government findings (2012) that also identified intellectual engagement, motivation, and identification with STEM pursuits as critical for persistence in STEM majors. These behavioral correlates, with impulsivity in particular, may deserve more consideration among faculty, STEM programs, as well as secondary and postsecondary institutions when supporting struggling students in STEM.

## Acknowledgements

The authors would like to thank Berk Moss (1945-2018) for guiding recruitment and retention procedures when working with schools and teachers.

